# Distribution of myogenic stem cell activator, hepatocyte growth factor, in skeletal muscle extracellular matrix and effect of short-term disuse and reloading

**DOI:** 10.1101/2025.03.14.643206

**Authors:** So Kuwakado, Alaa Elgaabari, Kahona Zushi, Sakiho Tanaka, Miyumi Seki, Ryuki Kaneko, Yuji Matsuyoshi, Takahiro Suzuki, Kenichi Kawaguchi, Yasuharu Nakashima, Ryuichi Tatsumi

## Abstract

Hepatocyte growth factor (HGF) is a key myogenic stem cells (satellite cells) activator, that resides in the extracellular matrix (ECM). However, HGF distribution in the ECM varies depending on the muscle fiber type. Furthermore, aging impedes the binding of HGF to its receptors due to nitration by peroxynitrite (ONOO^-^). Though oxidative stress increases rapidly during muscle disuse atrophy, satellite cells are rapidly activated upon reloading. In this study, we investigated the distribution of HGF in the ECM in various muscle fiber types, and examined nitration of HGF in disuse and reloading models. Immunofluorescence staining was performed on the soleus (Sol), plantaris (Pla), and gastrocnemius (Gas) muscles of 10-week-old mice. Three mice were used to assess HGF distribution, while 12 mice, divided into control, disuse, and reloading groups were used for qualitative evaluation of nitrated HGF (nitroHGF). In Sol muscle, type IIa and IIx muscle fibers exhibited higher HGF distribution in the ECM (70.7±1.9% and 63.9±2.6%, respectively) than type I fibers (28.0±1.7%; *p*<0.001). In Pla and Gas muscle, type IIa (65.4±2.3% and 70.3±2.2%, respectively) and type IIx fibers (55.3±1.8% and 61.4±3.2%, respectively) had significantly higher HGF distribution in the ECM than type IIb fibers (13.1±1.8% and 5.9±1.6%; *p*<0.001, respectively). The amount of nitroHGF increased in the disuse group compared to that in the control group but decreased in the reloading group compared to that in the disuse group. This preferential HGF distribution around type IIa and IIx muscle fibers indicates a distinct mechanism for satellite cell activation, differing from the satellite cell-rich environment associated with type I fibers and the lower HGF association with type IIb fibers. Disuse-induced HGF nitration may inhibit satellite cell activation. Reloading likely triggers mechanisms that counteract nitration, enabling satellite cell reactivation in young muscle.

## Introduction

In the hypertrophy and atrophy of skeletal muscles, apart from the balance between muscle protein synthesis (MPS) and muscle protein breakdown (MPB) rates [1–3], muscle stem cells (satellite cells) repair damaged areas of the muscle and cause hypertrophy, thus providing new myonuclear cells. Satellite cells reside in the niche between the sarcolemma and the extracellular matrix (ECM) of the muscle fiber, and most of them are dormant [4,5]. Satellite cells are activated by generating an activation signal cascade through Ca channels that sense changes in skeletal muscles during exercise as mechanical stimulation and undergo proliferation, differentiation, and fusion to become myotubes, which fuse to muscle injury sites, providing new myonuclear [5,6]. Although satellite cells are more abundant around slow-twitch muscle fibers, and myonuclear number increases predominantly in type I muscle fibers [7,8], resistance training preferentially induces increase in the number of myonuclear cells and hypertrophy in type II muscle fibers compared to type I muscle fibers [9]. In contrast, in clinical studies of older people who underwent habitual resistance training, type I muscle fibers showed increased myonuclear numbers, but type II muscle fibers did not [10]. This suggests that type II muscle fibers may have a different mechanism for controlling satellite cell activation than that of type I muscle fibers. As a possible candidate, we focused on the distribution of hepatocyte growth factor (HGF), a myogenic stem cell activator, in the ECM.

The activation signaling cascade of satellite cells includes a process in which HGF binds to the cell membrane receptor c-met [6]. HGF is abundant in the ECM of uninjured muscle tissue [11,12] and is released rapidly from extracellular tethering when subjected to mechanical stretching in living muscle [13–16]. The internal layer of the ECM surrounding muscle fibers has two primary constituents, collagen type IV and laminin-2, which vary in distribution as a function of muscle fiber type [17]. Qualitative evaluation has shown that the distribution of HGF in the ECM is biased depending on the muscle fiber type [11,18]. Therefore, the primary objective of the present study was to quantitatively evaluate the distribution of HGF in the ECM surrounding different muscle fiber types. We achieved this by referencing the measurement method of the activation rates of neuronal nitric oxide (NO) synthase (nNOS) [19].

Furthermore, our research group discovered that HGF in the ECM was nitrated with aging and lost its ability to bind to the c-met [18,20]. Aging induces a chronic and gradual increase in oxidative stress, which contributes to sarcopenia [21–24]. Nitrotyrosine is a promising biomarker of oxidative stress caused by the nitration of protein-bound tyrosine residues by ONOO^-^ formed from the major messenger NO and superoxide (O ^-^) [25]. When the tyrosine residue (Y198, Y250), present on the binding surface of HGF with the receptor c-Met is nitrated by ONOO^-^, it becomes nitroHGF, which shows poor binding possibly due to a change in polarity, resulting in the failure to activate muscle satellite cells and repair of muscle damage sites [18,20]. Elgaabari et al. (2024) reported that nitroHGF accumulates in the ECM of skeletal muscles of mice and rats with age [18]. An effect of aging on skeletal muscle tissue is the inactivation of satellite cells. Englund et al. (2020) reported that young mice treated with tamoxifen to inhibit satellite cells showed slower growth in the long term [26]. Thus, if the involvement of satellite cells induced by HGF in the ECM is important in human fast-twitch fibers, the accumulation of nitroHGF in the ECM is thought to be an essential etiology of sarcopenia.

Similar to the increase in oxidative stress associated with aging, oxidative stress also increases rapidly in disuse-induced muscle atrophy [27–30]. Therefore, increased nitroHGF likely inhibits satellite cell activation, even in disuse muscle atrophy. However, satellite cells are rapidly activated upon reloading after disuse [31]. Based on this, the second objective of this study was to investigate how disuse and reloading affect nitration of HGF and contribute to the characteristics of each muscle fiber type. This was assessed qualitatively using immunofluorescence staining.

## Materials and Methods

### Ethical approval

Animal interventions were conducted in strict accordance with the recommendations of the Guidelines for Proper Conduct of Animal Experiments published by the Science Council of Japan and with ethical approval from the Kyushu University Institutional Review Board (A22-082-2).

### Animals

Three C57BL6/J mice (male, 9 weeks) were ordered from KBT Oriental Co Ltd, bred, and housed at 23 ± 2 °C and 55 ± 10% humidity on a 12 h light/dark cycle (lights on at 8 a.m.) with free access to regular food (CRF-1, Oriental Yeast Co., Ltd.; Tokyo, Japan) and water. After a week of recovery, the animals were euthanized and tissues were collected. The animals were euthanized in a non-fasted state with ad libitum access to food and moisture. The Sol, Pla, and Gas muscles of the left lower hind limb were dissected from each mouse, oriented in tissue OCT compound, and frozen in isopentane cooled with liquid nitrogen.

Twelve C57BL6/J mice (male, 9 weeks old) were obtained from KBT Oriental Co., Ltd. and bred under the same conditions as described above. After one week of recovery, mice were randomly assigned to three groups: control group, disuse group, and reloading group, with four animals per group. After each group’s intervention period ended, the animals were euthanized, and tissues were collected. After measuring body weight, the tibialis anterior muscle (TA), extensor digitorum longus muscle (EDL), Sol, and Gas of the right lower hind limb were individually dissected from each mouse, and the muscle weights were measured. The Sol, Pla, and Gas muscles of the left lower hind limb were dissected as a lump from each mouse, oriented in tissue OCT compound, and frozen in isopentane cooled with liquid nitrogen.

### Hindlimb unloading

We used a disuse model of rodent hindlimb unloading [32] in which a strip of adhesive tape was attached to the tail of the mice. The tape was passed through a hole in the cage lid attached to a clip on the top of the cage. The animals could move around the cage, but their hind limbs could not touch the cage floor. Mice in the control group were unrestricted and allowed to move freely for 5 days. The disuse group mice were adapted to the disuse model and kept in a state of hindlimb unloading for 5 days. The reloading group mice were first adapted to the disuse model for 5 days and then allowed to spend the next 3 days free after removing the strip and releasing from hindlimb unloading. All mice were housed in a temperature-controlled room on a 12:12-h light-dark cycle with food pellets and transport agar (ORIENTAL YEAST Co., Ltd.) provided ad libitum.

### Direct-immunofluorescence microscopy

Some sets of serial cryo-sections (about 10-μm thickness) were prepared using a Leica CM1950 cryostat (Nussloch, Germany) and examined for muscle fiber type, distribution rate of HGF in ECM, short diameter of muscle fibers, or qualitative evaluation of HGF and nitroHGF by direct-immunofluorescence microscopy.

Muscle cross-sections were fixed with hot PBS and steam for 5 min and blocked with sterile donkey-serum solution (containing 2% normal serum, 1% BSA, 0.1% cold fish skin gelatin, 0.05% Tween 20, 0.01% avidin, 100 mM glycine, and 0.05% sodium azide in PBS, pH 7.2) for 1 h at 25°C, and stained with HiLyte Fluor™ 647-labeled anti-MyHC 1 monoclonal antibody (1:100 dilution), Alexa Fluor™ 350-labeled anti-MyHC 2A monoclonal antibody (1:100 dilution), HiLyte Fluorescein 500-labeled anti-MyHC 2X monoclonal antibody (1:200 dilution) and Alexa Fluor™ 594-labeled anti-MyHC 2B monoclonal antibody (1:200 dilution) overnight at 4°C. Then, other set of serial cryo-sections were fixed with hot PBS and steam for 5 min and blocked with sterile donkey-serum solution for 1 h at 25°C, and stained with HiLyte Fluor™ 647-labeled anti-MyHC 1 monoclonal antibody (1:100 dilution), Alexa Fluor™ 350-labeled anti-MyHC 2A monoclonal antibody (1:100 dilution), Alexa Fluor™ 594-labeled anti-MyHC 2B monoclonal antibody (1:200 dilution) and anti-laminin monoclonal antibody (Sigma-Aldrich, Burlington, MA, USA, L9393; 1:50 dilution) overnight at 4°C, and were then probed with a TRITC-conjugated goat anti-rabbit IgG secondary antibody (Thermo Fisher, Waltham, MA, USA, 1515529; 1:250 dilution) for 1 h at RT. In addition, another set of serial cryo-sections were fixed with citric acid buffer heated in microwave for 5 min, treated with TrueBlack (Biotium, Fremont, CA, USA) to suppress autofluorescence due to lipofuscin for 5 min and blocked with 3% BSA for 1 h at 25 °C before incubation overnight at 4 °C in Alexa Fluor 594-labeled anti-HGF monoclonal antibody (1:50 dilution in the sterile solution containing 1% BSA, 0.1% cold fish skin gelatin, 0.05% Tween 20, and 0.05% sodium azide in PBS) and anti-Laminin monoclonal antibody (Sigma-Aldrich, L9393; 1:50 dilution) overnight at 4°C, and were then probed with a TRITC-conjugated goat anti-rabbit IgG secondary antibody (Thermo Fisher, 1515529; 1:250 dilution) for 1 h at RT.

In next experiment, one set cross-sections were fixed with hot PBS and steam for 5 min and blocked with sterile donkey-serum solution for 1 h at 25°C, and stained with HiLyte Fluor™ 647-labeled anti-MyHC 1 monoclonal antibody (1:100 dilution), Alexa Fluor™ 350-labeled anti-MyHC 2A monoclonal antibody (1:100 dilution), HiLyte Fluorescein 500-labeled anti-MyHC 2X monoclonal antibody (1:200 dilution) and Alexa Fluor™ 594-labeled anti-MyHC 2B monoclonal antibody (1:200 dilution) overnight at 4°C. One other set of serial cryosections was fixed with citric acid buffer heated in a microwave for 5 min, treated with TrueBlack (Biotium) to suppress autofluorescence due to lipofuscin for 5 min, and blocked with 3% BSA for 1 h at 25°C before incubation overnight at 4°C in Alexa Fluor 594-labeled anti-HGF monoclonal antibody (1:50 dilution). Another set of serial cryo-sections were stained with Fluorescein 500-labeled anti-nitrated Y198 HGF monoclonal antibody (1:50 dilution) overnight at 4°C [18,33].

Sections were mounted in VECTASHIELD Antifade Mounting Medium (Vector Lab., Burlingame, CA, USA) and observed under a THUNDER Imager Live Cell (Leica microsystems). Micrographs were analyzed with morphometric measurements (ImageJ software) to calculate the length of BL rebelled with anti-laminin or anti-HGF antibody, and short diameter of muscle fibers, surrounding 10 muscle fibers randomly selected for each muscle fiber type within the section.

### Statistical analyses

Student’s t-tests were employed for statistical analysis of experimental results using Microsoft Excel X. Data are represented as mean ± standard error for 3 or 4 mice per group. The level of significance was set to p<0.05, and a statistically significant difference between the two groups at p<0.05 and at p<0.01 is indicated throughout by (*) and (**). Results are representative of independent experiments.

## Results

### Distribution of HGF in ECM at each muscle fiber type

The muscle fiber type proportion was obtained by quadruple immunostaining for the different myosin heavy chain (MyHC) isoforms (n=3). In soleus (Sol) muscle, type I muscle fibers accounted for 29.0%, type IIa muscle fibers were 67.0%, and type IIx muscle fibers were 4.0%. Plantaris (Pla) and gastrocnemius (Gas) muscles contained almost no type I muscle fibers, and type IIa muscle fibers were less abundant than in Sol muscle (22.5% and 7.4%). Type IIx muscle fibers were more abundant in Pla muscle (19.5%) and increased in number but only in a small proportion in the Gas muscle (5.9%). However, type IIb muscle fibers were absent in Sol muscle but accounted for 58.0% and 86.2% of Pla and Gas muscles (Fig 1a). Thus, in rodents used in animal experiments, slow-twitch muscles are abundant in type I and type IIa muscle fibers and contain a few type IIx muscle fibers.

**Fig 1.**
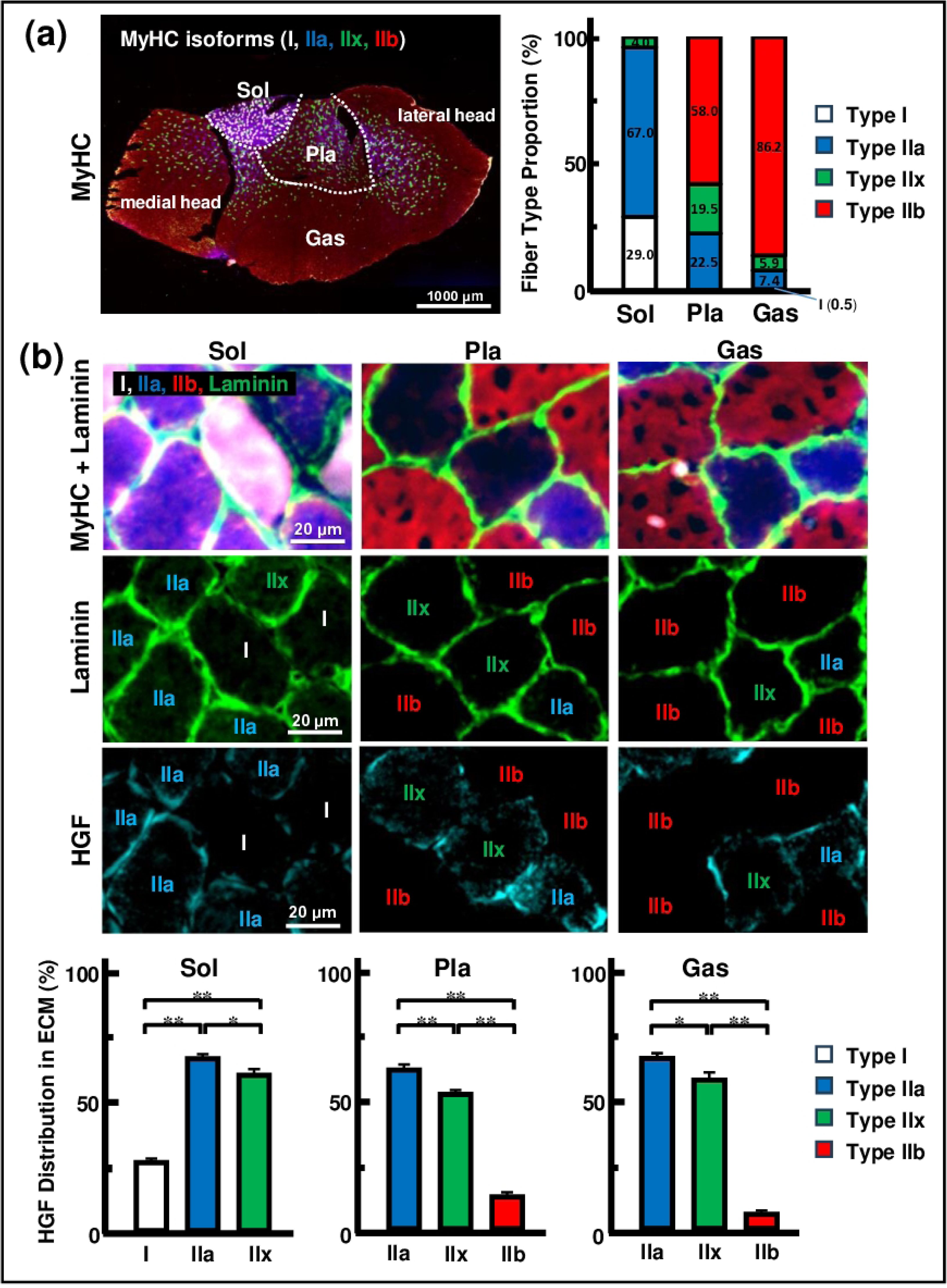
The distribution of HGF in ECM around each muscle fiber types. [a] Proportion of muscle fiber types in the Sol, Pla, and Gas muscles (n=3). In rodents used in animal experiments, slow-twitch muscles are abundant in type I and type IIa muscle fibers and contain few type IIx muscle fibers. In contrast, fast-twitch muscles are abundant in type IIb muscle fibers, contain some type IIa and IIx muscle fibers, and have few type IIx muscle fibers. [b] HGF distribution in the ECM of each muscle fiber type in the Sol, Pla, and Gas muscles (n=3). The length of the ECM surrounding each muscle fiber was determined using an anti-laminin antibody. Total HGF length in the ECM was determined using an anti-HGF antibody. In the Sol, IIa and IIx muscle fibers had a higher HGF distribution in the ECM than I muscle fibers (p<0.01). In the Pla and Gas muscles, IIa and IIx muscle fibers had a higher HGF distribution in the ECM than IIb muscle fibers (p<0.01). MyHC: Myosin heavy chain, Sol: Soleus muscle, Pla: Plantaris muscle, Gas: Gastrocnemius muscle, HGF: Hepatocyte growth factor. Statistically significant difference between the two groups at p<0.05 is indicated throughout by (*), and p<0.01 is indicated throughout by (**).

Next, we created co-stained images of MyHC I, IIa, IIb, and laminin, which are abundant proteins in the ECM, and of HGF and laminin from serial sections to visualize the distribution of HGF in the ECM surrounding each muscle fiber (Fig 1b). Ten muscle fibers were randomly selected for each muscle fiber type from the stained images of each muscle, and the length of ECM surrounding each muscle fiber labeled with anti-laminin antibody or the HGF distribution on ECM labeled with anti-HGF antibody was measured using Image J, and the distribution rate was calculated (n=3). Total HGF length in the ECM was determined using an anti-HGF antibody. In Sol muscle, type IIa, IIx muscle fibers (70.7 ± 1.9%, 63.9 ± 2.6%) have higher HGF distribution in ECM than type I muscle fibers (28.0 ± 1.7%; p<0.001), and type IIa muscle fibers have higher HGF distribution in ECM than type IIx muscle fibers (p=0.037). In Pla muscle, IIa, IIx muscle fibers (65.4 ± 2.3%, 55.3 ± 1.8%) have higher HGF distribution in ECM than IIb muscle fibers (13.1 ± 1.8%; p<0.001), and IIa muscle fibers have higher HGF distribution in ECM than IIx muscle fibers (p=0.001). Additionally, in Gas muscle, IIa, IIx muscle fibers (70.3 ± 2.2%, 61.4 ± 3.2%) have higher HGF distribution in ECM than IIb muscle fibers (6.0 ± 1.6%; p<0.001), and IIa muscle fibers have higher HGF distribution in ECM than IIx muscle fibers (p=0.025).

### Changes in short diameter and proportion of muscle fiber type due to disuse and reloading model

In the second experiment, 12 wild-type mice were divided into three groups and housed for 5 or 8 days: a control group, a disuse group, and a reloading group (Fig 2a). In the disuse model, muscle atrophy is most rapid in type I muscle fibers with abundant mitochondria producing O_2_ ^-^ [27–30]. Therefore, ONOO^-^ seems to be formed by O_2_ ^-^ reacting with NO more easily in slow-twitch muscles such as Sol [18,20]. In addition, type IIa and IIx muscle fibers with abundant HGF in the ECM constituted the majority of Sol. Therefore, in this experiment, we focused on the sol muscle in the disuse and reloading models. Short-term disuse did not induce significant atrophy in muscle weight/body weight ratio of Sol muscle (n=4) among control (1.00 ± 0.03), disuse (0.97 ± 0.06) and reloading group (0.98 ± 0.03) (Fig 2b). Ten muscle fibers were randomly selected for each muscle fiber type from the stained images of the Sol muscles in each group, and the Minimal Feret Diameter of each muscle fiber, as the short diameter, was measured using ImageJ (n=4). Type I muscle fibers was longer in the control group (34.8 ± 1.0 µm) than in the disuse (31.1 ± 0.7 µm; p<0.001) and reloading groups (30.7 ± 0.6 µm; p<0.001). Type IIa muscle fibers had no difference among the control (32.9 ± 0.9 µm), disuse (31.8 ± 0.8 µm), and reloading groups (31.6 ± 0.8 µm). Type IIx muscle fibers were longer in the disuse group (35.6 ± 1.2 µm) than in the control group (32.5 ± 0.9 µm; p=0.047), but not in the reloading group (33.8 ± 0.8 µm). (Fig 2c). In the proportion of muscle fiber types (n=4), type I muscle fibers had no difference among control (35.5 ± 4.7%), disuse (30.3 ± 0.5%), and reloading groups (33.7 ± 1.7%). Type IIa muscle fibers were smaller in the disuse group (58.0 ± 2.3%) than in the reloading group (49.5 ± 3.1%; p=0.074) but not in the control group (56.6 ± 5.2%). Type IIx muscle fibers were larger in the reloading group (16.8 ± 2.1%) than in the control group (7.9 ± 0.6%; p=0.006), but not in the disuse group (11.7 ± 2.3%) (Fig 2d).

**Fig 2.**
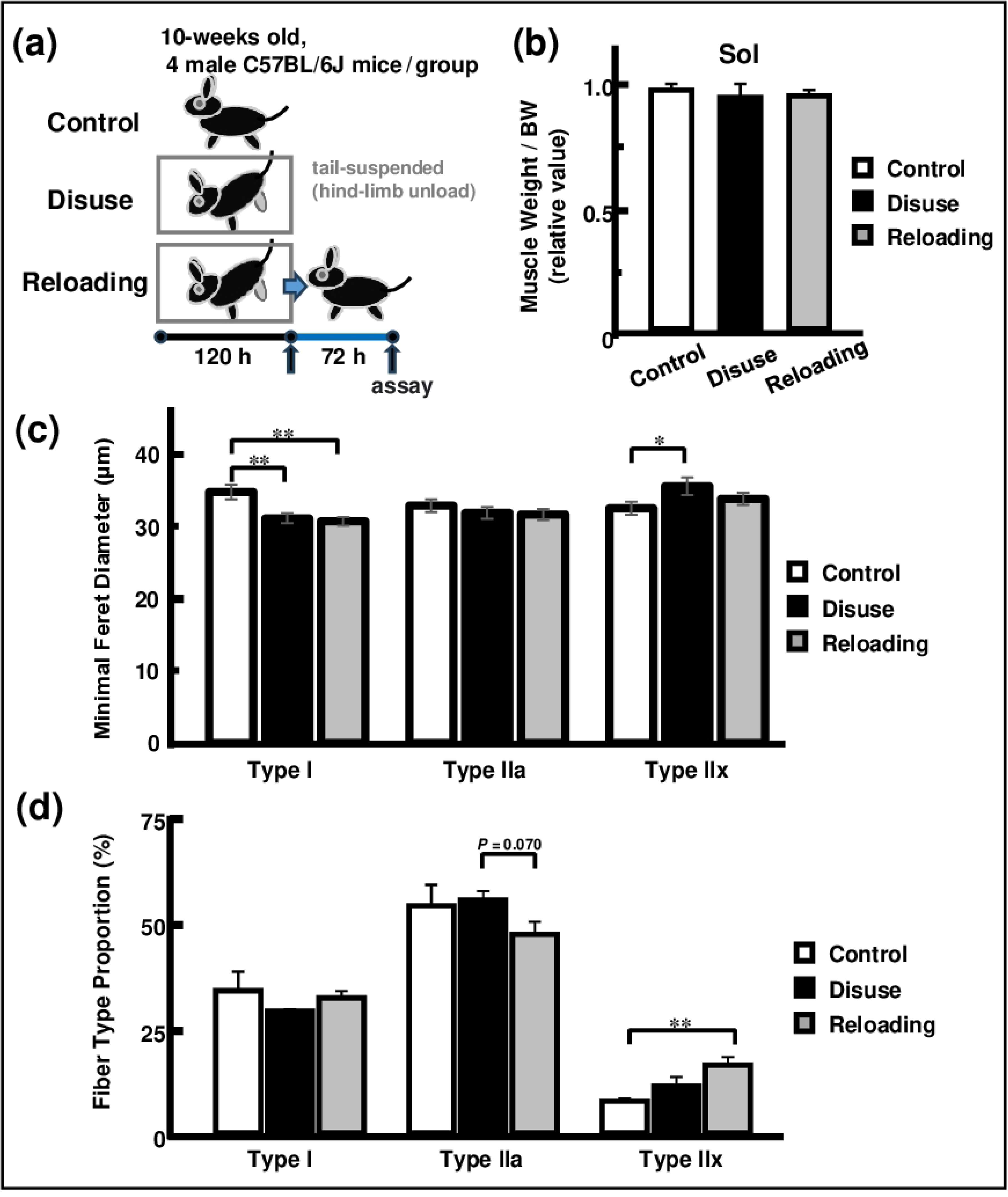
Muscle fiber types in short-term disuse and short-term reloading model. [a] Intervention schedule for the experiment. The mice in the control group (n=4) spent 5 days freely. The mice in the disuse group (n=4) were subjected to hind limb unloading for 5 days. The mice in the reloading group (n=4) were adapted to hind limb unloading for 5 days and allowed to spend the next 3 days freely. [b] Relative muscle weight to body weight in the control, disuse, and reloading groups (n=4). The control, disuse, and reloading groups showed no significant differences. [c] Minimal Feret diameter of each muscle fiber in the Sol (n=4). Type I muscle fibers decreased in the disuse and reload groups compared to the control group (p<0.05). Type IIx muscle fibers increased in the disuse group compared to those in the control group (p<0.05). [d] Proportions of muscle fiber types in the Sol for the control, disuse, and reloading groups (n=4). Type IIa muscle fibers tended to decrease in the reloading group compared with the disuse group (p=0.074). Type IIx muscle fibers increased in the reloading group compared to those in the control group (p<0.05). BW: Body weight. Statistically significant difference between the two groups at p<0.05 is indicated throughout by (*), and p<0.01 is indicated throughout by (**).

### Signal intensity of HGF and nitroHGF due to disuse and reloading model

In the qualitative evaluation based on antibody signal intensity, the amount of nitro-HGF increased in the disuse group compared to that in the control group. In contrast, the amount of nitro-HGF decreased in the reloading group compared to that in the disuse group (Fig 3a).

**Fig 3.**
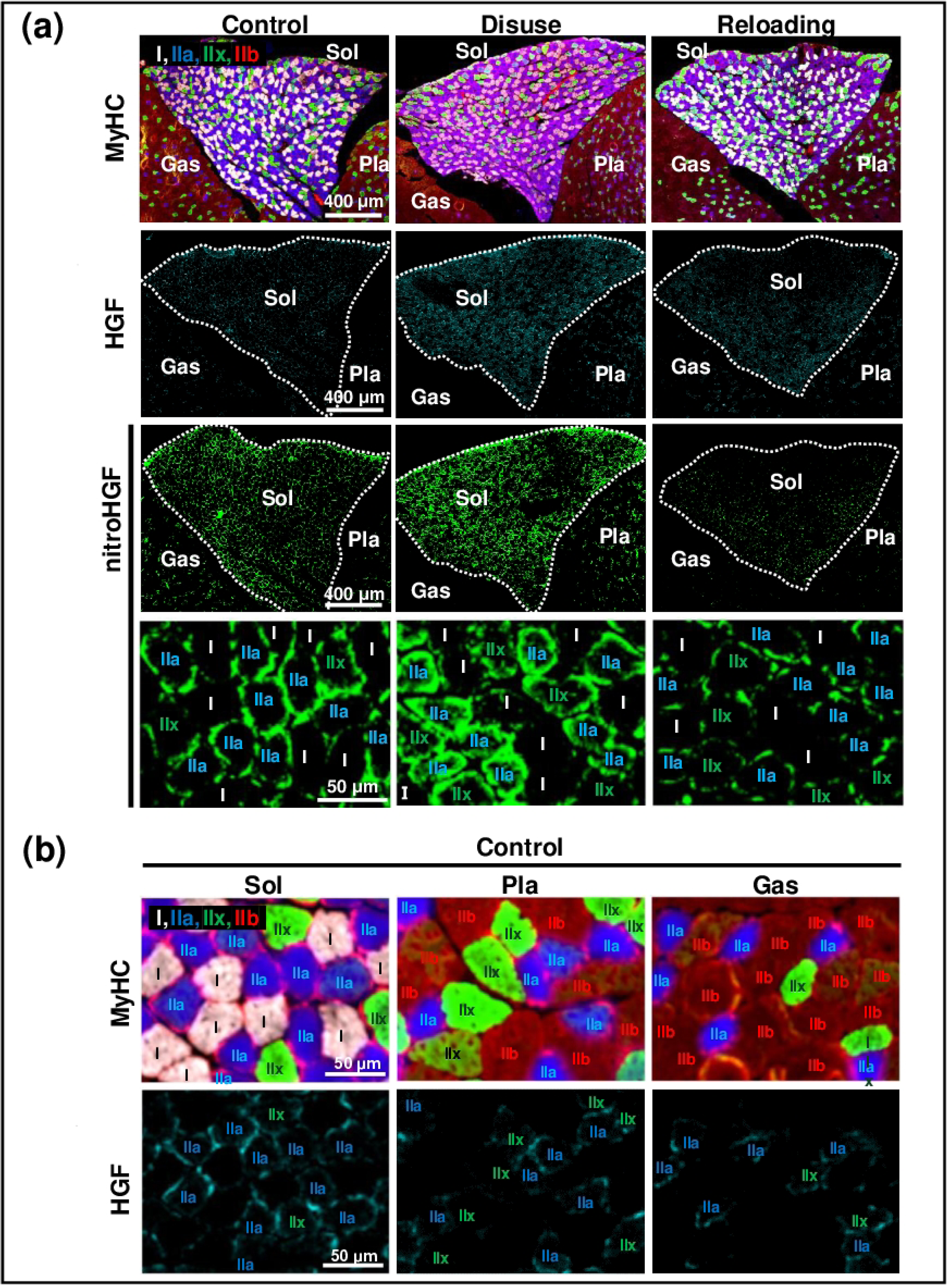
Nitration and increase in HGF levels in short-term disuse and reloading models by immunofluorescence staining. [a] Qualitative evaluation of immunofluorescence staining with anti-HGF and anti-nitro-HGF antibodies. NitroHGF in the ECM increased in the disuse group compared to that in the control group and decreased in the reloading group composed of the disuse group. In addition, HGF in the ECM increased in the disuse group compared to the control group and decreased in the reloading group composed to disuse group. [b] Qualitative evaluation of immunofluorescence staining with anti-HGF antibody in the control group. The amount of HGF around type IIa and IIx muscle fibers in the Sol was higher than that in the Pla and Gas muscles.

Additionally, in the qualitative evaluation based on antibody signal intensity, the amount of HGF increased in the disuse group compared to the control group and decreased in the reloading group compared to the disuse group (Fig 3a). To determine whether immobility due to the disuse model itself was the cause or whether increased oxidative stress was the cause, we focused on HGF around type IIa and type IIx muscle fibers in each muscle from the stained images of the control group, since slow-twitch muscles such as the Sol muscle have abundant type I muscle fibers that contain a large number of mitochondria, which physiologically generate oxidative stress [34]. In the control group, the Sol muscle had more HGF in the ECM around type IIa and IIx muscle fibers than the Pla and Gas muscles (Fig 3b).

## Discussion

The distribution of HGF in the ECM has not received as much attention as its distribution in satellite cells. This may be because type I muscle fibers, around which satellite cells are abundant [7,8], are the largest in slow-twitch muscles, whereas type IIb muscle fibers are the largest and most abundant in fast-twitch muscles (S1 and S2 Figs). However, since type I and IIa muscle fibers account for a high proportion of large mammals, such as humans, and type IIb is almost absent [35], medical research on humans should focus on HGF in the ECM surrounding type IIa and IIx muscle fibers as a characteristic of fast-twitch fibers and the activation of muscle satellite cells via the HGF/c-met pathway. Around IIb muscle fibers, satellite cells are less prevalent, and HGF is not widely distributed in the ECM. Therefore, type IIb muscle fibers may have mechanisms to uniquely enhance the self-repair capacity of muscle fibers [36] and MPS metabolism [1–3], which may not be mediated by satellite cell activation via the HGF/c-met pathway.

Many rodent studies that address myonuclear numbers have not focused on the distinction between type II muscle fibers, and few studies have shown clear benefits of the abundance of HGF in the ECM, which is characteristic of type IIa and IIx muscle fibers. A study on myonuclear number in humans reported that resistance exercise [35], especially unaccustomed exercise habits [37], increased the myonuclear number in type II muscle fibers more than in type I muscle fibers. In addition, a study focusing on mouse soleus muscles reported that type IIa and IIx muscle fibers hypertrophied more rapidly than type I muscle fibers in long-term reloading experiments after long-term disuse [38]. However, as reported in other experimental studies of disuse muscle atrophy and spinal cord injury models [35,39], we observed a transition to the muscle fiber type that reduces durability in the slow-twitch muscle Sol in our short-term disuse and reloading model. The difference in the average size of type IIa and IIx muscle fibers was significant, but it disappeared due to the transition between muscle fiber types (S3 Fig). This transition may make it difficult to evaluate the characteristic hypertrophy for each muscle fiber type. Further research to address the timing of muscle fiber type transition and innervation back to the original type [35] is needed to understand the unique relationship between satellite cells and each muscle fiber type.

In the short-term disuse model used in the second experiment, the muscle weight/body weight ratio of the Sol showed no significant change. However, oxidative stress was generated to the extent that the short diameter of the type I muscle fibers was significantly reduced. Therefore, a certain amount of O_2_^-^ was generated in our short-term disuse experiment, possibly resulting in the formation of ONOO^-^. Qualitative evaluation showed that nitroHGF increased in the skeletal muscle ECM of the disuse group. This suggests that similar to aging, the activation of satellite cells is suppressed when exercise is resumed in disuse muscle atrophy. In contrast, nitroHGF decreased in the reloading group, and the release of HGF from the ECM did not seem to be inhibited. Although this is not a direct proof, this series of results can be explained by the reduction or decomposition of nitroHGF with reloading. However, since HGF reacts rapidly after being released by exercise from an ECM that has been anchored in advance and is not synthesized step-by-step [13–16], the decomposition of nitroHGF and resynthesis of HGF are unlikely. Since a rapid reaction occurs in muscle fibers due to ONOO^-^ in the early stages of exercise [40] and exercise is repeated, researchers can reasonably assume that a mechanism reducing nitration modification may exist in muscle fibers. The mechanism, thereby assisting in the reactivation of satellite cells present in muscle fibers, may be like cytosolic enzymes such as CK [41] or LDH [42], which are released outside the cell when the sarcolemma is destroyed by exercise. This is a possible reason why the rapid hypertrophy of slow muscles is not suppressed when loading is resumed after disuse in young mice and rats [31].

Although disuse muscle atrophy is rapid [27–30], especially in young people, satellite cells are activated more rapidly upon reloading and contribute to the recovery of skeletal muscle mass than usual [31]. In the qualitative evaluation, the HGF levels in the disuse group were higher than those in the control group. This may be a mechanism that activates satellite cells more than usual, which is the reason for the rapid hypertrophy during reloading [31]. In the control group, even around the same type IIa and type IIx muscle fibers, the ECM of the Sol muscle appeared to contain more HGF than the Pla and Gas muscles. Slow-twitch muscles such as Sol are mostly made up of muscle fibers with many mitochondria, such as type I and type IIa muscle fibers [43]. Therefore, a larger amount of oxidative stress is generated than fast-twitch muscles such as Pla and Gas on a normal basis [34]. Although the detailed expression mechanism remains unknown, oxidative stress likely promotes HGF expression. In reloading after disuse, Sol muscle promoted the transition to type IIx muscle fibers, and this transition to type II muscle fibers is also induced by regular resistance exercise [9]. Few reports address resistance exercise. However, similar to aerobic exercise, an increase in oxidative stress has been reported [44]. In addition, Li, et al. (2024) reported the oxidative stress from endurance exercise promotes the transition of fast-twitch to slow-twitch muscle fibers, which contrasts with our findings [45]. It remains unclear whether aerobic or resistance exercise generates stronger oxidative stress. However, disuse muscle atrophy is associated with a substantial increase in reactive oxygen species, which causes rapid muscle atrophy [30]. Therefore, oxidative stress caused by disuse or exercise may play a role in modulating muscle fiber type transitions.

Our experiment has six limitations: (1) The study was conducted on mice, and similar reactions in humans remain unguaranteed. Using human muscle tissue samples, confirming that HGF in the ECM is abundant around type IIa and IIx muscle fibers is necessary. (2) A clear characterization of HGF abundance in the ECM is required. Increasing the myonuclear number and administering tamoxifen must clearly distinguish between type IIa, IIx, and type IIb muscle fibers. (3) We used only nitrotyrosine as an oxidative stress marker, and the generation of reactive oxygen species was indirectly indicated by the atrophy of type I muscle fibers. A more detailed evaluation would require the use of other oxidative stress markers. (4) Other more accurate methods may exist for quantifying HGF levels in the ECM surrounding each muscle fiber. A higher magnification microscope might enable us to determine which side of the ECM, labeled with the anti-laminin antibody, HGF is. (5) More quantitative methods are required to determine the degree of nitro-HGF. HGF is present in extremely low amounts in the skeletal muscle tissue, making it difficult to quantify by western blotting. However, a quantitative evaluation is necessary for a clear evaluation. (6) More detailed verification, such as reverse transcription polymerase chain reaction (PCR) is required regarding the increased expression of HGF in response to oxidative stress and the conditions for muscle fiber-type transition.

In conclusion, HGF may be abundant in the ECM around type IIa and IIx muscle fibers, providing new myonuclear cells and rendering them susceptible to muscle repair and hypertrophy by satellite cells via the HGF/c-Met pathway. This seems to be a different control mechanism from that of type I muscle fibers, which are surrounded by abundant satellite cells and also distinguishs type IIa and IIx from type IIb muscle fibers. In disuse muscle atrophy, nitroHGF increases in response to oxidative stress. Satellite cells can be reactivated by a reduction mechanism for nitration during reloading, which has a rapid effect on type IIa and IIx muscle fibers in young individuals.

## Authors’ Contributions

S.K. performed the experiments, analyzed the data, interpreted the experimental results, prepared the figures, and drafted the manuscript; A.E., S.T., M.S., K.Z., R.K., and Y.M. supported the experimental techniques; S.K. and K.K. edited and revised the manuscript; T.S., Y.N., and R.T. approved the final version of the manuscript.

## Acknowledgments

Special thanks to Ms. Akiko Sato (Kyushu University, Japan) for the technical assistance with animal care. The authors thank Muscle & Meat Sci. Lab, Department of Animal and Marine Bioresource Sciences, Graduate School of Agriculture for the chance of the experiments. The authors thank the Center for Advanced Equipment and Educational Support, Faculty of Agriculture, Kyushu University for the use of the fluorescence microscope. This work was supported by Grants-in-Aid for Scientific Research (B) from the Japan Society for the Promotion of Science (JSPS KAKENHI, Grant Numbers 21H02347 and 24K01911) (all to R.T.). This research was supported in part by grant funds from the Uehara Memorial Foundation and the Ito Foundation (to R.T.). The funders had no role in the study design, data collection and analysis, decision to publish, or preparation of the manuscript and did not provide support in the form of salaries for any author. We would like to thank Editage (www.editage.jp) for English language editing.

## Data availability

All data generated and analyzed in this study are included in the paper and the Supporting Information.

## Supporting Information

**S1 Fig. Minimal Feret diameter of each muscle fiber in the first experiment (n=3).** In Sol muscle, type I muscle fibers (34.3 ± 0.9 µm) are larger than type IIa (30.6 ± 1.2 µm; p=0.015) and IIx (30.2 ± 0.7 µm; p<0.001). In Pla and Gas muscles, type IIb muscle fibers (42.3 ± 1.4 µm and 45.0 ± 1.7 µm) are larger than type IIa (27.1 ± 1.2 µm and 23.2 ± 1.1 µm; p<0.001) and IIx (34.6 ± 1.3 µm and 29.0 ± 1.2 µm; p<0.001).

MyHC, myosin heavy chain; Sol: Soleus muscle; Pla: Plantaris muscle; Gas: Gastrocnemius muscle; HGF: Hepatocyte growth factor. Statistically significant differences between the two groups at p<0.05, as indicated by (*), and p<0.01 is indicated throughout by (**).

**S2 Fig. Muscle fiber type proportions of Sol, Pla, and Gas muscles in the control group in the second experiment (n=4).**

In Sol muscle, type I muscle fibers accounted for 33.2%, type IIa muscle fibers were 59.2%, and type IIx muscle fibers were 7.6%. In Pla muscle, type IIa muscle fibers accounted for 10.0%, type IIx muscle fibers for 13.5%, and type IIb muscle fibers for 76.5%. In Gas muscle, type I muscle fibers accounted for 0.6%, type IIa muscle fibers for 3.7%, type IIx muscle fibers for 4.5%, and type IIb muscle fibers for 91.2%.

**S3 Fig. Minimum Feret diameter of each muscle fiber in Sol (n=4).**

In the control group, type I (34.8 ± 1.0 µm), IIa (32.9 ± 0.9 µm), and IIx muscle fibers (32.5 ± 0.9 µm) showed no differences in size. In disuse group, type IIx muscle fibers (35.6 ± 1.2 µm) were larger than type I (31.1 ± 0.7 µm; p=0.002) and IIa (31.9 ± 0.8 µm; p=0.015). In reloading group, type IIx muscle fibers (33.8 ± 0.8 µm) were larger than type I muscle fibers (30.7 ± 0.6 µm; p<0.001) but only tended to be than type IIa muscle fibers (31.6 ± 0.8 µm; p=0.056).

**S4 Fig. Qualitative evaluation of immunofluorescence staining with anti-HGF and anti-nitro-HGF antibodies in other individuals.**

NitroHGF in the ECM increased in the disuse group compared to that in the control group and decreased in the reloading group composed of the disuse group. In addition, HGF in the ECM increased in the disuse group compared to the control group and decreased in the reloading group composed of the disuse group.

**S5 Fig. Qualitative evaluation of immunofluorescence staining with anti-HGF antibody in the control group.**

The amount of HGF around type IIa and IIx muscle fibers in the Sol was higher than that in the Pla and Gas muscles.

**S1 Table. Proportion of muscle fiber types in the Sol, Pla, and Gas muscles (n=3).**

This is the table of data in Fig 1a.

**S2 Table. HGF distribution in the ECM of each muscle fiber type in the Sol, Pla, and Gas muscles (n=3).**

This is the table of data in Fig 1b.

**S3 Table. Relative muscle weight to body weight in the control, disuse, and reloading groups (n=4).**

This is the table of data in Fig 2b.

**S4 Table. Minimal Feret diameter of each muscle fiber in the Sol (n=4).**

This is the table of data in Fig 2c.

**S5 Table. Proportions of muscle fiber types in the Sol for the control, disuse, and reloading groups (n=4).**

This is the table of data in Fig 2d.

